# Strain-specific response to pharmacological cold mimicking in aging mice

**DOI:** 10.64898/2026.01.08.698532

**Authors:** Roshan Lal, Ojas Tikoo, Neha Soni, Arka Bhattacharya, Kanthi Kiran Kondepudi, Kanwaljit Chopra, Mahendra Bishnoi

## Abstract

**Background:** Aging impairs thermogenic and metabolic flexibility, increasing metabolic disease risk. We examined six aging male mouse strains for responses to chronic topical menthol, a pharmacological cold mimetic.

**Material and methods:** Male (C57BL/6J, A/J, BALB/c, C3H/hej, DBA/2J, and FVB/NJ) mice were treated with 4g/kg of 10% menthol weight per volume cream once per day or vehicle as control using finger application for 15 days. After last application, core body temperature measurement, BAT thermography, Nesting behaviour, and metabolic tolerance test were performed, and gene expression was analysed in BAT.

**Results:** Menthol improved core temperature recovery via thermogenesis and enhanced insulin sensitivity across all strains. However, effects on BAT mass, nesting behaviour, and gluconeogenesis were strain-dependent.

**Conclusion:** Collectively, these findings highlight that genetic background modulates metabolic responses to cold mimetics, informing future precision strategies for age-related metabolic disorders.

## 1. Introduction

Aging is a biological phenomenon characterized by progressive decline in physiological tissue function. It is associated with loss in metabolic adaptability, increasing the susceptibility to metabolic disorders [1]. Dysregulation of energy homeostasis during aging contributes to unbalanced energy flux, resulting in excess fat storage in white adipose tissue (WAT). Over time, WAT hypertrophy leads to the initiation of metabolic disorders like type 2 diabetes, hyperlipidaemia & other obesity complications [2, 3]. To counterbalance excess energy stored, activation of energy expenditure/utilization pathways becomes essential to achieve balanced energy flux. Cold stimuli increase energy expenditure in response to maintain homoeothermy, which becomes impaired due to aging [4]. Cold exposure primarily acts through skin’s sensory neurons [5] that relay temperature changes to the hypothalamus through Transient Receptor Potential (TRP) channels [3, 6]. Hypothalamus, activates thermoregulatory responses through Brown adipose tissue (BAT) and skeletal muscle [3, 7]. Elevating basal energy expenditure through pharmacological cold mimicking is an intervention strategy for fat loss. Menthol is a TRPM8 (innocuous cold, <16-25°C) agonist, and its topical application activates thermogenesis in assorted ways [8] to tilt crucial energy storage to expenditure. Action of chronic topical menthol treatment is well studied for its role in transient increment of basal thermogenesis through TRPM8 signalling [9]. However, due to phenotypic factors like age, sex, diet, physical activity, etc., menthol’s pharmacological cold action across multiple genotypes as a preventive anti-obesity therapeutic remains obscure. So, to better investigate such changes, here, we have taken aging male mice of six different strains and studied their response to pharmacological cold mimic, topical menthol.

## 2. Methods

### 2.1 Animals

Male mice (22-23 weeks old) of six strains C57BL/6J (24-26g), A/J (24-26g), BALB/c (30-32g), C3H/hej, (30-32g), DBA/2J (28-30g), FVB/NJ (30-32g) were obtained from Advanced Centre for Treatment, Research and Education in Cancer (ACTREC), Mumbai. Animals were housed in Animal Experimentation Facility (AEF) at National Agri-Food and Biomanufacturing Institute (NABI), Mohali, India, under standard conditions: 23 ± 2°C, 55 ± 5% humidity, and a 12-h light/dark cycle with ad libitum access to food and water. All procedures complied with the Committee for the Purpose of Control and Supervision of Experiments on Animals (CCSEA) guidelines and were approved by Institutional Animal Ethics Committee (IAEC) of NABI (Approval No. NABI/2039/CPCSEA/IAEC/2022/14).

Following acclimatization, mice were randomized into two groups (n = 3–5 per strain per group). (1) Vehicle (2) Menthol (4 gm/kg 10 % w/v menthol). Topical menthol and vehicle were applied across the interscapular to sacral region once a day over two weeks (15 times in total) [9].

### 2.2 Core-body temperature (CBT) Monitoring

CBT was measured with a rectal probe (Orchid Scientific and Innovative India Pvt. Ltd) at 0 (before menthol application), 30, 60, 120, and 180 min post-menthol application [9]

### 2.3 Infrared thermography

Thermogenic activity of the intrascapular BAT (iBAT) was assessed by surface area temperature monitoring through IR thermography. Briefly, pictures of mice were taken at 120 mins post-menthol application using FLIR T530 thermal camera (emissivity 0.98). Two standardized regions of interest, ROI-1 (interscapular region) and ROI-2 (nuchal/neck region), are chosen for calculation using FLIR tools app [9].

### 2.4 Metabolic tolerance tests

Pyruvate tolerance test (PTT) (1□g/kg sodium pyruvate (Cat. no. P5280), 12□h fasting), and insulin tolerance test (ITT) (0.75□IU/kg human insulin (Cat. no. I9278), 4□h fasting) conducted and blood glucose was monitored using Freestyle glucometer using Freestyle glucose strips at 0, 15, 30, 45, 60, 90, 120 min [9].

Cold tolerance test (CTT) was performed in 16 h fasted mice housed at 4°C for 4h with free access to water. Rectal temperature was measured every hour during cold exposure using a rectal probe. All metabolic tests were performed after 15-day menthol application [10].

### 2.5 Nesting behaviour

Nesting behaviour was assessed after 15 days of topical menthol application. Mice were individually housed without enrichment, and a nestlet (3 gm) was placed. Nest formation was evaluated as described previously [9].

### 2.6 RT-qPCR analysis

Briefly, total RNA from Liver, BAT, and hypothalamus was extracted using the Trizol-based method and reverse transcribed into cDNA using RevertAid cDNA synthesis Kit (#K1622) [11], and qPCR was performed (CFX96) with iTaq SYBR Green (#1725121). Cycling conditions: 95°C for 2 min; 40 cycles of 95°C for 5 s and 60°C for 30 s. Relative gene expression was calculated via the 2−ΔΔCT method [12], normalized to TBP. Primers are listed in **Table S1**.

### 2.7 Statistical analysis

Graphpad Prism 8 software (Graphpad, California, United States) was used for statistical analysis. Data are expressed as mean ± SEM and analysed using two-tailed unpaired t-test (with Welch’s correction); p < 0.05 was considered statistically significant. Correlation analysis was done with Pearson’s coefficients (corrplot, R), and significant values are shown. Modified custom summary data dashboard was created using Python.

## 3 Results

### 3.1 Topical menthol stimulates differential phenotypic and genotypic sensitivity to cold-like behaviour with BAT activation

We used six independent mouse strains: C57BL/6J, A/J, BALB/c, DBA/2J, C3H/HeJ, and FVB/NJ **(Fig. 1A)** that genetically belong to distinct groups; group 1 (A/J, BALB/c, C3H/HeJ), group 2 (FVB/NJ), group 4 (C57BL/6J) & group 6 (DBA/2J) as per Petkov et. al [13]. Topical application of menthol (15 days) led to enhanced nest-building behaviour, indicating increased cold sensitivity and behavioural thermoregulation across all strains, with significant response in C57BL/6J, DBA/2J, and FVB/NJ (p < 0.05) mice **(Fig. 1E)**, Consistent with adaptive cold responses. Further, menthol application increases BAT (primary effector tissue in heat production) tissue mass significantly **(Fig. 1F)** in C57BL/6J (∼15.5%, p < 0.05), DBA/2J (∼16.4%, p < 0.05), and FVB/NJ mice (∼19.6%, p < 0.05) along with higher interscapular surface temperature **(Fig 1D)** in C57BL/6, C3H/HeJ, & FVB/NJ (p<0.05). These changes correlated with upregulated mitochondrial thermogenic markers (*UCP1* and *NRF2*, p<0.05) expression (**Fig. 1H)**. Notably, TRPC5 expression was increased in hypothalamus **(Fig. 1H)** of C57BL/6J, BALB/c, C3H/HeJ, and FVB/NJ (p < 0.05), suggesting enhanced central cold-sensing mechanisms.

**Figure 1.**
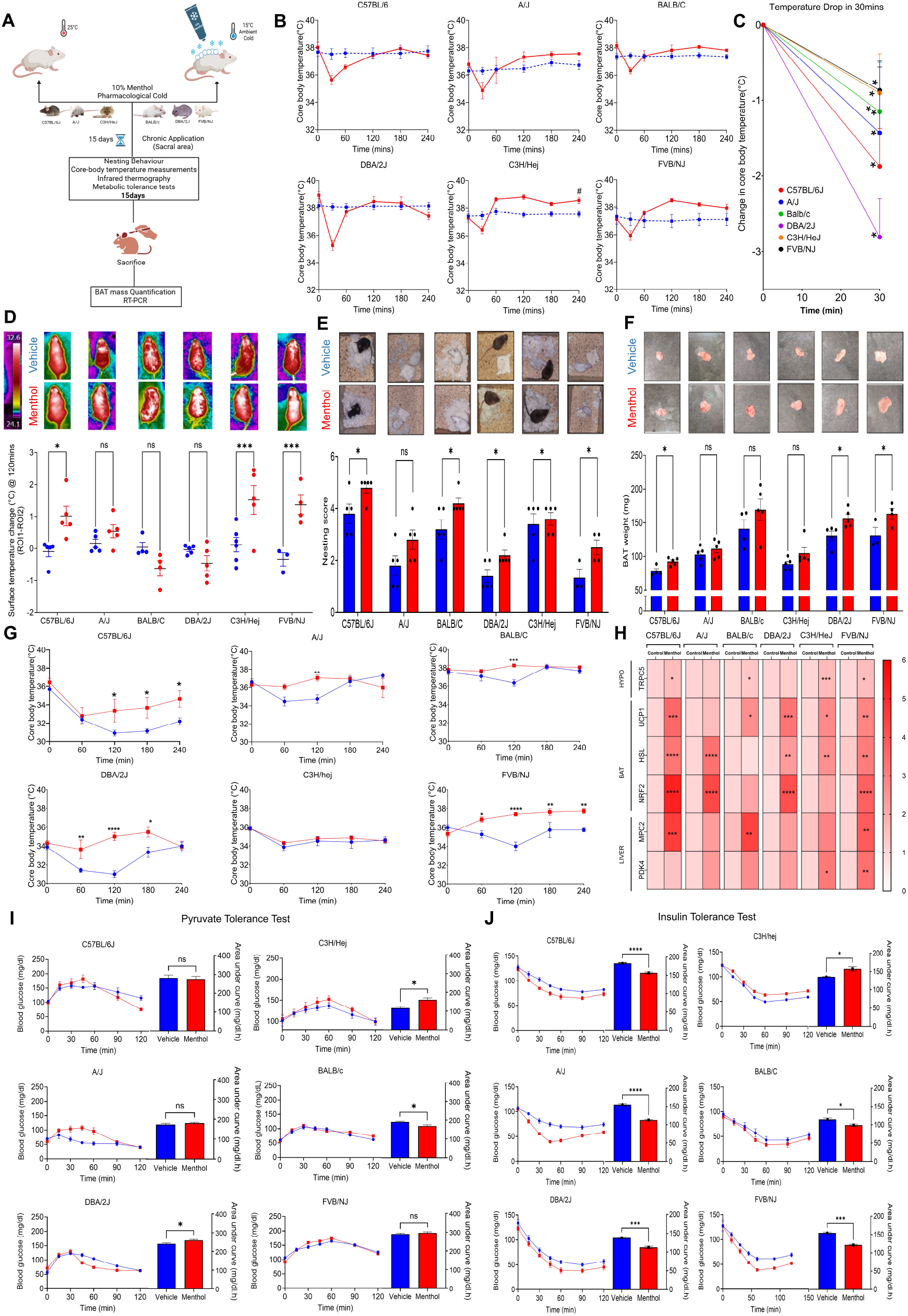

### 3.1 Topical menthol actuates differential cold sensing, thermostasis, and metabolic expenditure

CBT measurements in vehicle vs menthol application demonstrated globally an initial reduction in CBT by approximately 3–8% across all six strains of menthol-treated mice, followed by restoration. This shows loss in body heat as an effect of menthol’s TRPM8 stimulation **(Fig. 1B, p<0.05)**. Besides C57BL/6J, menthol group relayed an early thermoregulatory response and restored thermostasis in 60 min **(Fig. 1C, p<0.05)**. CTT in vehicle vs 15-day menthol application revealed that menthol-treated mice returned to baseline core temperatures within 60 minutes, indicating effective cold acclimatization through improved thermal homeostasis in all the strains **(Fig. 1G)**. Further, ITT, C57BL/6J, A/J, DBA/2J, and FVB/NJ (p < 0.05) mice treated with menthol showed a significant decrease in blood glucose concentration, 45–60 minutes post-injection, indicating improved insulin responsiveness **(Fig. 1J, p<0.05)**. PTT revealed increased glucose production from pyruvate in C57BL/6J, A/J, and DBA/2J strains following 15-day menthol treatment, consistent with enhanced hepatic gluconeogenesis **(Fig. 1I, p<0.05)**, genes *MPC2 & PDK4* **(Fig. 1H, p<0.05)**.

### 3.3 Correlation of cold-sensing, thermogenic pathways & metabolic status under pharmacological Cold Mimicking

Strain-specific correlation analysis showed pharmacological cold mimicking linking to *TRPC5, UCP1 & HSL* (p<0.5), which is what drives global flow of energy expenditure to menthol’s effect. Strain-specific co-relation analysis represents the afferent thermogenic effect that drives the global non-shivering BAT thermogenesis **(Fig. 2A)**. Co-relation analysis shows which pathway pharmacological cold acclimatization affects based on genotypic variations. Beyond *UCP1-HSL*’s positively co-related pair to menthol’s action, it shows differences in genetically insulin-tolerant FVB/NJ’s significant co-relation with increased insulin sensitivity upon menthol application.

**Figure 2.**
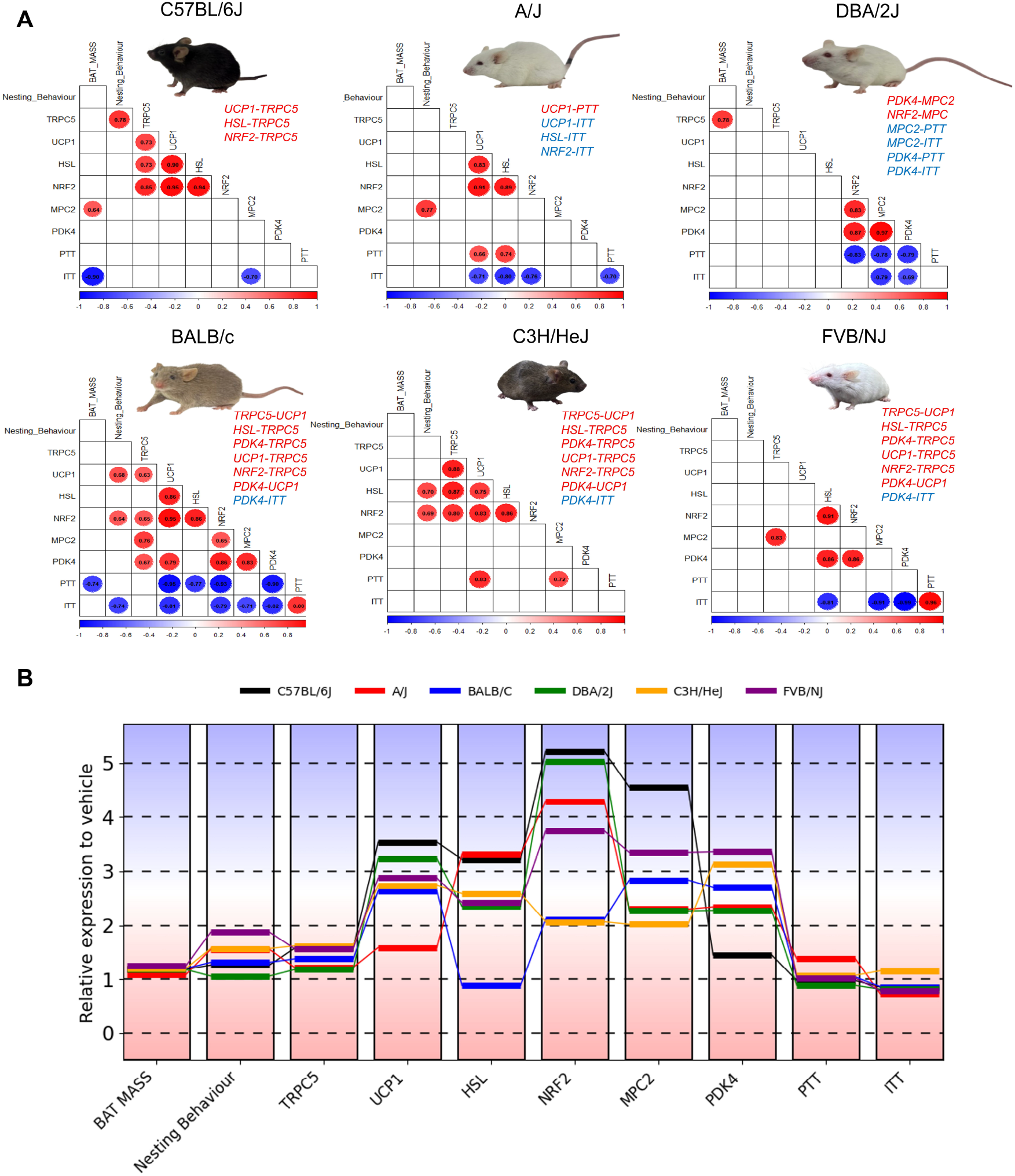

## 4. Discussion

We investigated how genotypically distinct aging mouse strains respond to pharmacological cold mimicking via recurrent TRPM8 activation. Menthol treatment reproduced key features of environmental cold acclimatization, including enhanced nest-building behaviour and elevated interscapular BAT surface temperature, consistent with increased thermoregulation and energy expenditure [9, 14]. These behavioural and physiological changes were validated by significant increases in BAT mass and upregulation of mitochondrial thermogenic genes (UCP1, NRF2), supporting improved thermostasis under cold-like conditions [2, 3, 9]. Further, metabolic profiling revealed that menthol enhanced systemic insulin sensitivity and promoted cold-induced metabolic flexibility [2, 15]. Strain-specific differences in response likely reflect genetic background, relative cold sensitivity, TRPM8 saturation, and tolerance to thermal variation. Importantly, pharmacological cold differs mechanistically from environmental cold: TRPM8-mediated perception induces internal heat generation with subsequent heat loss to a neutral environment, whereas environmental cold drives thermogenesis to counter external heat sink– mediated loss. Lastly, correlation analyses across six strains identified significant associations between hypothalamic TRPC5 expression and TRPM8 agonist response, linking central cold sensing to peripheral metabolic pathways in liver (MPC2, PDK4) and BAT (UCP1, HSL, NRF2), particularly in genetic groups 1, 2, and 4 [13, 16]. Collectively, these findings demonstrate that cold sensitivity and adaptive thermogenesis are strongly genotype-dependent **(Fig. 2B)** and that pharmacological cold mimicking offers a targeted strategy to activate TRP channels and enhance energy expenditure in aging mice.

## 5. Conclusion

Overall, our study concludes the role of genotype in the action of pharmacological cold mimicking using topical menthol application. This points to the significance of personalized/group therapeutics based on specific genotypes.

## Data Availability

Source data are provided with this paper. Any other data reported in this paper are available from the corresponding author upon reasonable request.

## Acknowledgement

The authors acknowledge ICMR, GoI, for providing fellowship to Roshan Lal and Ojas Tikoo. Authors also thank CSIR, UGC, GoI and National Agri-food and Biomanufacturing Institute (NABI) for providing fellowship to Neha Soni and Arka Bhattacharya. We acknowledge DBT e-Library Consortium (DeLCON) for literature support. The funders had no role in study design, data collection and analysis, decision to publish, or preparation of the manuscript.

## Author’s Contribution

MB conceived the idea and designed the experiments. R.L, O.T, and N.S performed the experiments. M.B, R.L, and O.T analysed the data. O.T and M.B wrote the manuscript. O.T and A.B made the figures. M.B, K.C, and K.K helped in the manuscript writing and data presentation.

---

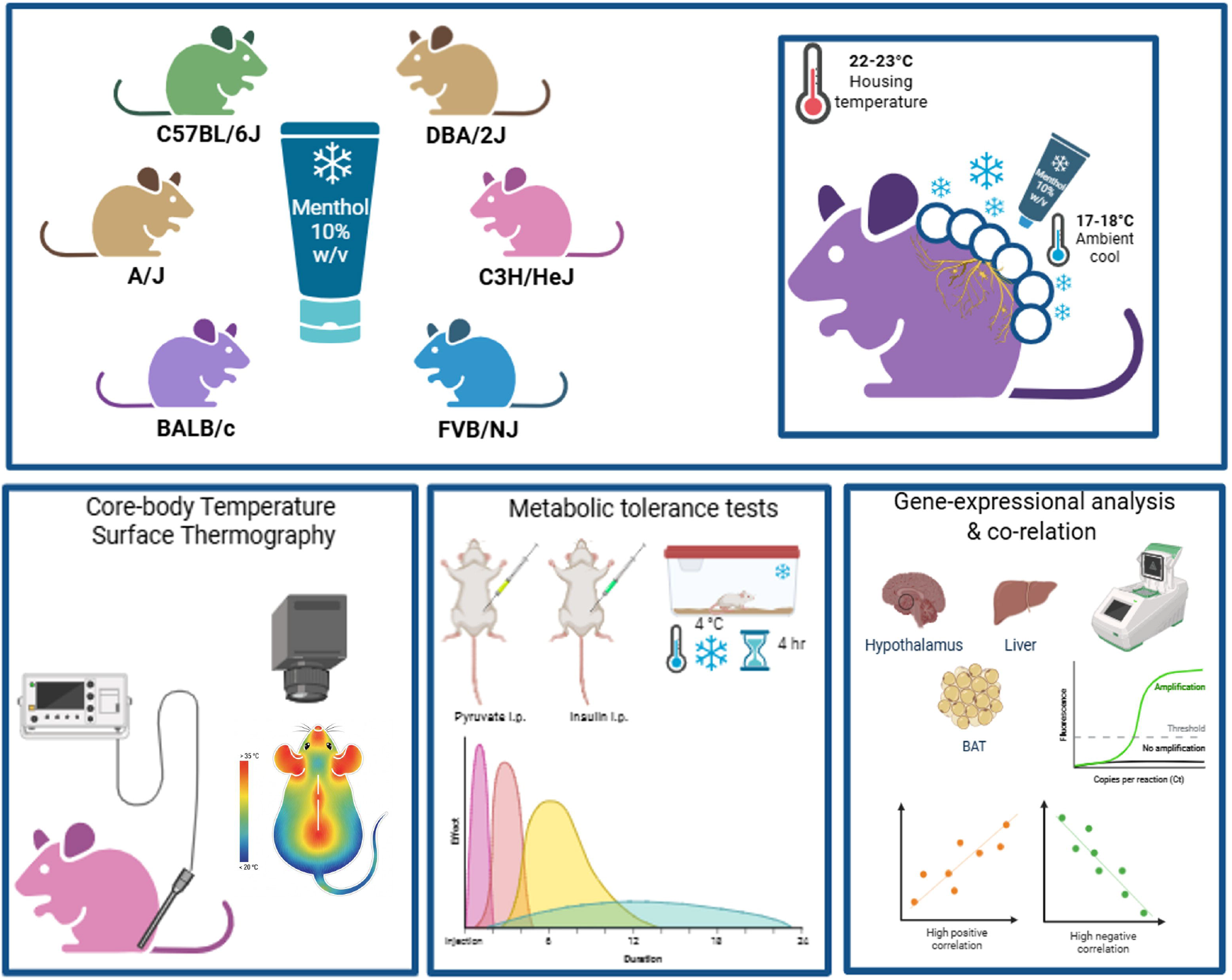

